# A short isoform of the UNC-6/Netrin receptor UNC-5 is required for growth cone polarity and robust growth cone protrusion in *Caenorhabditis elegans*

**DOI:** 10.1101/2023.05.02.539117

**Authors:** Snehal S. Mahadik, Erik A. Lundquist

**Affiliations:** Program in Molecular, Cellular, and Developmental Biology Department of Molecular Biosciences The University of Kansas 1200 Sunnyside Avenue 5049 Haworth Hall Lawrence, KS 66045

## Abstract

UNC-6/Netrin is a conserved bi-functional guidance cue which regulates dorsal-ventral axon guidance in *C. elegans*. In the Polarity/Protrusion model of UNC-6/Netrin mediated dorsal growth away from UNC-6/Netrin, The UNC-5 receptor first polarizes the VD growth cone such that filopodial protrusions are biased dorsally. Based on this polarity, the UNC-40/DCC receptor stimulates growth cone lamellipodial and filopodial protrusion dorsally. The UNC-5 receptor maintains dorsal polarity of protrusion, and inhibits growth cone protrusion ventrally, resulting in net dorsal growth cone advance. Work presented here demonstrates a novel role of a previously undescribed, conserved short isoform of UNC-5 (UNC-5B). UNC-5B lacks the cytoplasmic domains of UNC-5 long, including the DEATH domain, the UPA/DB domain, and most of the ZU5 domain. Mutations that specifically affect only the *unc-5* long isoforms were hypomorphic, suggesting a role of *unc-5B* short. A mutation specifically affecting *unc-5B* cause loss of dorsal polarity of protrusion and reduced growth cone filopodial protrusion, the opposite of *unc-5* long mutations. Transgenic expression of *unc-5B* partially rescued *unc-5* axon guidance defects, and resulted in large growth cones. Tyrosine 482 (Y482) in the cytoplasmic juxtamembrane region has been shown to be important for UNC-5 function, and is present in both UNC-5 long and UNC-5B short. Results reported here show that Y482 is required for the function of UNC-5 long and for some functions of UNC-5B short. Finally, genetic interactions with *unc-40* and *unc-6* suggest that UNC-5B short acts in parallel to UNC-6/Netrin to ensure robust growth cone lamellipodial protrusion. In sum, these results demonstrate a previously-undescribed role for the UNC-5B short isoform, which is required for dorsal polarity of growth cone filopodial protrusion and to stimulate growth cone protrusion, in contrast to the previously-described role of UNC-5 long in inhibiting growth cone protrusion.

## Introduction

Axons are guided to their targets in the developing nervous system by extracellular cues that provide guidance information. Receptors for these guidance cues on the growth cones of extending axons sense and respond to these guidance cues (GALLO AND LETOURNEAU 1999). In *C. elegans*, UNC-6/Netrin is a conserved, extracellular laminin-like guidance cue, and is expressed in ventral cells and axonal processes (ISHII *et al*. 1992; WADSWORTH *et al*. 1996). In response to UNC-some axons grow ventrally toward UNC-6, and some grow dorsally away from UNC-6 (HEDGECOCK *et al*. 1990).

In the Polarity/Protrusion model of VD growth cone outgrowth dorsally away from UNC-6 expression, UNC-6 first polarizes the growth cone such that filopodial protrusions are directed dorsally, and then regulates protrusion of the growth cone based on this polarity (NORRIS AND LUNDQUIST 2011; NORRIS *et al*. 2014; GUJAR *et al*. 2017; GUJAR *et al*. 2018). The Polarity/Protrusion model is consistent with vertebrate studies showing that floorplate Netrin is dispensable for commissural axon growth (DOMINICI *et al*. 2017; VARADARAJAN AND BUTLER 2017; YAMAUCHI *et al*. 2017; MORALES 2018), and is not consistent with classical ventral gradient models (BOYER AND GUPTON 2018). The UNC-6 receptor UNC-5 (LEUNG-HAGESTEIJN 1992) is required for dorsal polarity of filopodial protrusion, and also inhibits protrusion laterally and ventrally in response to UNC-6 (NORRIS AND LUNDQUIST 2011; MAHADIK AND LUNDQUIST 2022). The UNC-6 receptor UNC-40 (CHAN *et al*. 1996) is required for robust dorsal filopodial protrusion in response to UNC-6 (NORRIS AND LUNDQUIST 2011). This balance of ventral inhibition of protrusion by UNC-5 and dorsal stimulation of protrusion by UNC-40 results in net dorsal growth away from UNC-6.

Previous studies implicate three distinct pathways utilized by UNC-5 to inhibit growth cone protrusion. The FMO flavin monooxygenases act downstream of UNC-5 to inhibit protrusion (GUJAR *et al*. 2017), possibly via oxidation and destabilization of F-actin, similar to the FMO molecule MICAL. UNC-33/CRMP acts downstream of UNC-5 to inhibit protrusion by preventing microtubule + end entry into growth cones, which is pro-protrusive (GUJAR *et al*. 2018). TOM-1/Tomosyn acts downstream of UNC-5 to inhibit protrusion (MAHADIK AND LUNDQUIST 2023), likely by inhibiting the formation of the SNARE complex and thus vesicle fusion. This might prevent plasma membrane addition to the growth cone needed to extension, or prevent vesicles with pro-protrusive factors (*e.g.* Arp2/3) from being delivered to the growth cone plasma membrane.

The *unc-5* locus is predicted to encode for multiple isoforms (KILLEEN *et al*. 2002). Multiple isoforms differ at the 5’ end of the gene, resulting in UNC-5 molecules with different complements of the extracellular Immunoglobulin (Ig) and Thrombospondin (TSP) repeats. One isoform differs at the 3’ end of *unc-*5 gene, and is predicted to encode a molecule lacking most of the cytoplasmic domains, including the DEATH domain, the UPA/DB domain, and most of the ZU5 domain. Previous studies of the effects of UNC-5 on growth cone polarity and protrusion have utilized mutant alleles that affect all *unc-5* isoforms (*unc-5* strong loss-of-function mutants) (NORRIS AND LUNDQUIST 2011; NORRIS *et al*. 2014; GUJAR *et al*. 2018; GUJAR *et al*. 2019; MAHADIK AND LUNDQUIST 2022; Mahadik and Lundquist 2023).

Work reported here is an analysis of the role of the *unc-5B* short isoform in axon guidance and growth cone morphology. *unc-5* mutant alleles that affect only the long isoforms and not the short isoform were found to be hypomorphic for *unc-5* function compared to *unc-5* strong loss-of-function mutants: they displayed less severely uncoordinated locomotion and weaker VD/DD axon guidance defects. However, growth cone polarity and protrusion largely resembled strong *unc-5* mutants, with loss of dorsal polarization of filopodial protrusion and a larger growth cone area and longer filopodia compared to wild-type. Using CRISPR/Cas9 genome editing, an *unc-5* allele was generated that was predicted to affect only the *unc-5B* short isoform. This *unc-5B* mutant displayed weakly uncoordinated locomotion and weak VD/DD axon defects. *unc-5B* mutant growth cones lacked dorsal polarity of filopodial protrusion, indicating that UNC-5B is required for VD growth cone polarity. *unc-5B* mutant growth cones were also smaller and less protrusive than wild-type, indicating a possible pro-protrusive role for UNC-5B. *unc-5B* failed to complement a strong loss-of-function *unc-5* mutant for these phenotypes, but complemented *unc-5* hypomorphic mutants, consistent with the idea that *unc-5B* short encodes a distinct genetic function *unc-5* long isoforms. Transgenic expression of *unc-5B* rescued uncoordinated locomotion and ax on defects of *unc-5* mutants, and perturbed VD growth cone polarity and protrusion consistent with a pro-protrusive role.

In sum, work reported here is consistent with a role of the *unc-5B* short isoform in VD/DD axon guidance that is genetically separable from the function of the *unc-5* long isoforms. Both are required for VD growth cone dorsal polarity of protrusion, UNC-5 long isoforms are required to inhibit growth cone protrusion, and UNC-5B short isoform might have a pro-protrusive role. Vertebrate genomes encode four *Unc5* genes, *A, B, C, and D*, with *Unc5A* having a predominant role in axon guidance (ACKERMAN *et al*. 1997; LEONARDO *et al*. 1997; PRZYBORSKI *et al*. 1998). Human and other vertebrate *Unc5A* genes are predicted to encode a short isoform similar to *C. elegans unc-5B*, called *Unc5A X4*. Possibly, this isoform is acting similar to *unc-5B* in human nervous system development.

## Results

### UNC-5B encodes a short isoform lacking most cytoplasmic domains

The *unc-5* locus is predicted to encode at least seven isoforms, A-C, D1, D2, E and F (Wormbase WS245) (Figure 1A). Six of the seven isoforms (A, C-F) differ by exon usage at the 5’ end of the gene, resulting in distinct extracellular N-termini. *unc-5B* differs at the 3’ end from the other 5 isoforms, resulting in a truncated 3’ end (Figure 1A). The short isoform *unc-5B* is generated by skipping the 5’ splice donor site of intron 7 (second to the last intron) (Figure 1B). The open reading frame extends 44 bases into intron 7 before encountering a stop codon (Figure 1B), resulting in an additional 15 amino acid residues with no similarity to other known sequences. In RNA-seq of mixed stage animals, 242 reads contained the exon 7 to exon 8 splice, and 9 reads overlapped exon7/intron 7 or intron 7/exon 8 boundaries (Figure 1C). Neighboring introns 6 and 9 showed no reads overlapping the exon/intron boundaries, suggesting that the intron 7 reads are representative of *unc-5B.* These results indicate that *unc-5B* constituted 3.6% (9/251) of *unc-5* transcripts in mixed-stage animals.

**Figure 1.**
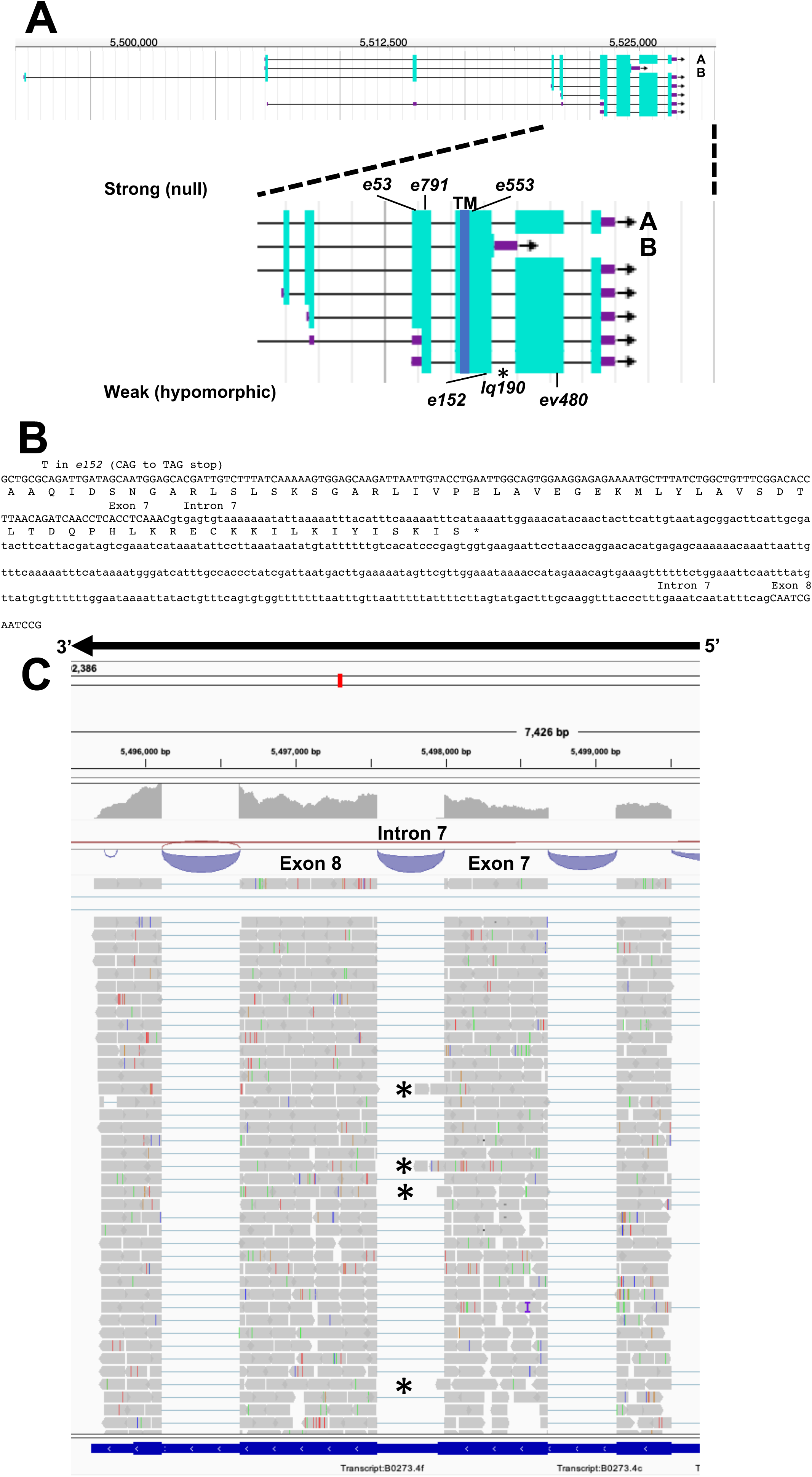
Gene structure of *unc-5* with isoforms and mutant alleles. (A) Gene structure of *unc-5* (Wormbase 287) showing five isoforms, A, C, D, F are long isoforms and B is short isoform. The positions of the strong alleles affecting all isoforms (*e53, e791I, and* e553) and the hypomorphic alleles affecting only the long isoforms (*ev480, op468*, and *e152*) are indicated on the expanded figure of the 3’ end on *unc-5*. E152 affects all isoforms but behaves as a hypomorph. The asterisk (*) indicates *unc-5(lq190)* a precise deletion of intron 7 predicted to abolish the *unc-5B* short isoform (B) The sequence of the exon7-intron 7-exon eight region. The C-terminus of the *unc-5B* short isoform is indicted in intron 7. (C) RNA seq reads in the exon7-intron 7-exon eight region from mixed stage wild-type animals is shown (from the Integrated Genome Browser). 5’ to 3’ is right to left. Reads representing the *unc-5B* short isoform are indicated with asterisks.

The UNC-5A long isoform encodes a molecule with two extracellular immunoglobulin (Ig) domains, two extracellular thrombospondin type I repeats (TSPI), a transmembrane domain, and cytoplasmic zona occludens I/UNC-5 domain (ZU5), UNC-5/PIDD/Ankyrin (UPA), and DEATH domains (Figure 2A). A tyrosine residue (Y482) that is phosphorylated and is important for functional UNC-5 (KILLEEN *et al*. 2002) is found in the intracellular juxtamembrane region (Figure 2A). The short isoform UNC-5B lacks the DEATH and UPA domains and most of the ZU5 domain, but retains Y482 (Figure 2A).

**Figure 2.**
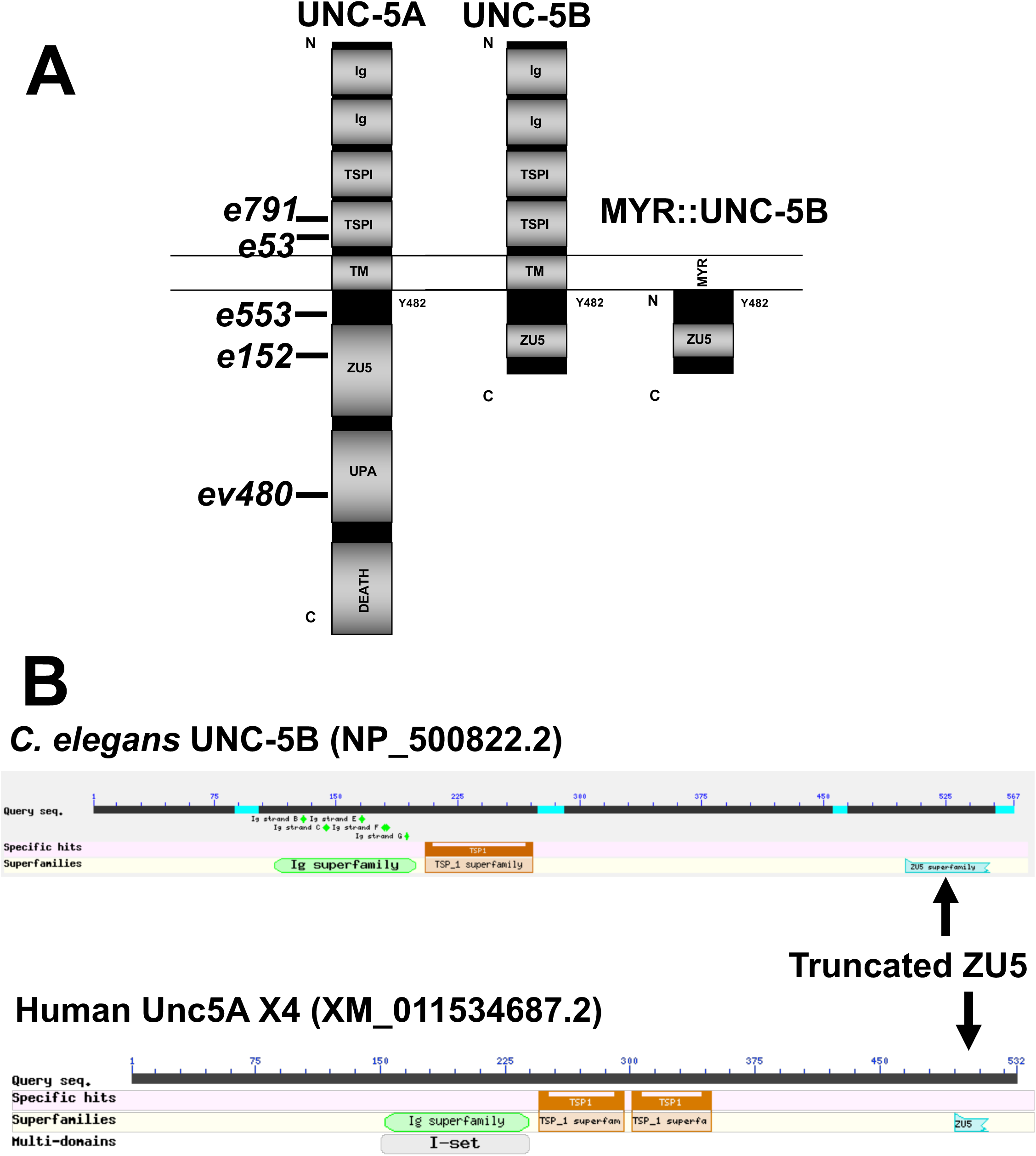
Domain structure of UNC-5A and UNC-5B. (A) The modular organization of UNC-5A and UNC-5B isoforms is shown. Domains include extracellular immunoglobulin domains (Ig) and thrombospondin domains (TSP), a transmembrane region (TM), and cytoplasmic ZU5, UPA, and DEATH domains. The position of Y482 in the cytoplasmic juxtamembrane region is indicated. A depiction of the myristoylated (MYR) version of the UNC-5B short cytoplasmic domain is shown. Positions of *unc-5* mutations are shown. (B) Graphic summaries of protein alignments from BLAST are shown for UNC-5B from *C. elegans* and UNC-5A X4 from *H. sapiens*. The truncated ZU5 domain at the C terminus of each is indicated.

Vertebrates, including humans, have four *unc5* genes, *A-C.* The human Unc5A locus encodes a truncated isoform similar to *C. elegans* UNC-5B, called Unc5A X4 (nucleotide XM_011534687.2, protein XP_011532989.1) (Figure 2B). Unc5A X4 also includes a portion of the ZU5 domain before ending with a stop codon (Figure 2B). *unc5A X4* isoforms are also predicted in other vertebrate species (*e.g.* mouse and *Xenopus*).

### Mutations predicted to affect only the UNC-5 long isoforms are hypomorphic

The *unc-5* isoforms represented by *unc-5A* are collectively referred to as *unc-5* long isoforms, and *unc-*5*B* the short isoform. The *unc-5* alleles *e791, e53, e553,* and *e152* introduce premature stop codons that affect all *unc-5* isoforms (Figures 1A and 2). The *ev480* and *op468* alleles affect only the *unc-5* long isoforms and do not affect the *unc-5B* short isoform (Figures 1A and 2). *ev480* introduces a premature stop codon, and *op468* is a 337-bp out-of-frame deletion. The *e791* and *e53* mutations predicted to affect all isoforms caused a strong uncoordinated locomotion (Unc) phenotype (Table 1). The long-isoform-specific mutations *ev480* and *op468* caused a weaker Unc phenotype (Table 1). The *e152* allele, which also is predicted to affect all isoforms, also caused a weaker Unc phenotype (Table 1). Based on axon guidance defects, *e152* is also a hypomorphic mutation (Table 1), similar to previous findings (KILLEEN *et al*. 2002).

**Table 1.** VD/DD axon guidance defects.

| Genotype<br>(n = 1600)<sup>1</sup> | Movement<br>(Unc<br>phenotype)<br><sub>2</sub> | % midline<br>crossing<br>defects<sup>3</sup> | % Total<br>axon<br>guidance<br>defects<sup>4</sup> | Comparison<br>* *p* < 0.001 |
| --- | --- | --- | --- | --- |
| *wild-type* | +++ | <1 | 1 |  |
| *unc-5(e791)* | + | 93 | 95 |  |
| *unc-5(e53)* | + | 96 | 97 |  |
| *unc-5(e152)* | ++ | 27* | 70* | *e791*, *e53*,<br>*ev480*, and<br>*op468* |
| *unc-5(ev480)* | ++ | 15* | 37* | *e791* and<br>*e53* |
| *unc-5(op468)* | ++ | 17* | 36* | *e791* and<br>*e53* |
| *unc-5A(+)*<sup>5</sup> | ++ | 10* | 12* | *wild-type* |
| *unc-5(e791); unc-5A(+)* | ++ | 9* | 13* | *e791* |
| *unc-5A(Y482F)*<sup>6</sup> | ++ | 6* | 28* | *wild-type* |
| *unc-5(e791); unc-5A(Y482F)* | ++ | 27* | 73* | *e791; unc-5A(+)* |
| *unc-5B(+)*<sup>5</sup> | ++ | 6* | 16* | *wild-type* |
| *unc-5(e791); unc-5B(+)* | ++ | 11 | 41* | *e791* and<br>*e791; unc-5A(+)* |
| *unc-5B(Y482F)*<sup>6</sup> | ++ | 28 | 56 | *wild-type* |
| *unc-5(e791); unc-5B(Y482F)* | + | 80 | 98 | *e791; unc-5B(+)* |
| *myr::unc-5A*<sup>7</sup> | ++ | 15* | 41* | *wild-type* |
| *myr::unc-5B*<sup>7</sup> | ++ | 9* | 10* | *wild-type* |
| *unc-5(lq190)* | +++ | <1 | 3* | *wild-type* |
| *unc-40(n324)* | ++ | 12 | 27 |  |
| *unc-40(n324); unc-5(lq190)* | ++ | 11 | 53* | *n324* |
| *unc-40(n324); unc-5(ev480)* | ++ | 61* | 86* | *n324* and<br>*ev480*<br>additive |
| *unc-6(ev400)* |  | 76 | 94 |  |
| *unc-5(lq190); unc-6(ev400)* |  | 89* | 100* | *ev400* |

Red highlights severe axon guidance defects ( $\leq 33\%$ )
Yellow highlights moderate axon guidance defects (34-66%)
Blue highlights mild axon guidance defects ( $\geq 67\%$ )
<sup>1</sup>100 animals were scored, and on average, 16 commissures consisting of the 19 VD/DD neurons were evident in wild-type animals.
<sup>2</sup>+++ equals wild-type locomotion; ++ equals slightly uncoordinated locomotion; + equals severely uncoordinated locomotion.
<sup>3</sup>The percent, rounded to the nearest percentile, of the 16 commissures evident in wild-type that did not cross the lateral midline in each genotype.
<sup>4</sup>The percent, rounded to the nearest percentile, of the 16 commissures evident in wild-type that has lateral midline crossing defects, pathfinding, ectopic branching, crossover and not able to reach to dorsal nerve cord.
<sup>5</sup>*unc-5A(+)* and *unc-5B(+)* represent transgenes harboring wild-type copies of each isoform (see Materials and Methods).
<sup>6</sup>*unc-5A(Y482F)* and *unc-5B(Y482F)* represent transgenes harboring copies of each isoform with the tyrosine 482 to phenylalanine mutation (see Materials and Methods).

VD/DD axon guidance defects were scored in these mutants. The 16 VD/DD commissures, composed of 19 axons, extend directly ventrally to dorsally (Figure 3A). *e791* and *e53* mutants displayed very severe VD/DD guidance defects, with between only 5-7% of axons extending dorsally past the lateral midline of the animal (Table 1 and Figure 3B). In contrast, *ev480, op468,* and *e152* displayed significantly less severe defects, with between 70-85% extending past the lateral midline (Table 1 and Figure 3C and D). Despite extending past the lateral midline, most axons in these mutants were still misguided as evidenced by lateral wandering (Figure 3C-D). These data indicate that *ev480, e152,* and *op468* are hypomorphic, incomplete loss-of-function mutations.

**Figure 3.**
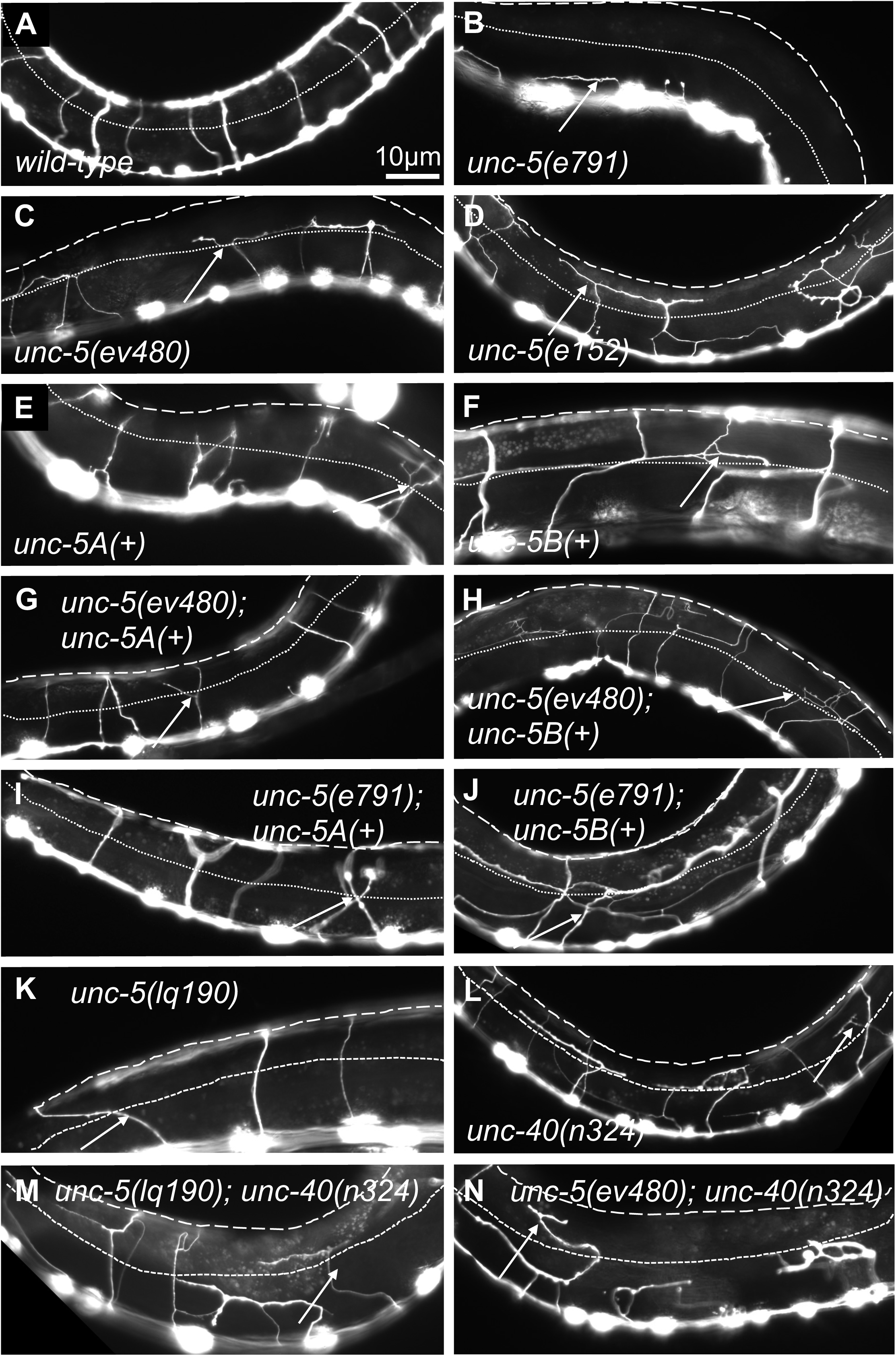
VD/DD axon guidance defects of mutants. Fluorescent micrographs of the *Punc-25::gfp* transgene *juIs76* expressed in the VD/DD neurons of L4 animals in different genotypes are shown. Dorsal is up and anterior left. The approximate lateral midline is indicated with a dotted white line, and the dorsal nerve cord by a dashed white line. White arrows indicate axon guidance defects in each genotype. Scale bar in (A) represents 10μm. Genotypes are indicated in each figure panel.

*ev480* and *op468* are predicted to affect only long *unc-5* isoforms. The hypomorphic *e152* mutation resides in exon 7 (Figure 1B), which is included in all six isoforms, yet *e152* has a hypomorphic phenotype (Table 1 and Figure 3). Exon 7 is the terminal exon in *unc-5B* short, and the *e152* mutation resides 183 nucleotides (61 codons) from the end of the predicted *unc-5B* reading frame. This is downstream of the transmembrane domain and Y482. Any truncated molecule made by translation of *e152* would include both (Figure 1). Most of the truncated ZU5 domain would also be included. Possibly, *unc-5B* short function is retained in *e152* by bypassing nonsense-mediated decay due to its location near the stop codon in the terminal exon. However, axon guidance defects in *e152* are significantly stronger than *ev480* and *op468* (Table 1), suggesting that *e152* might be having some effect on *unc-5* long isoforms, just not complete elimination of activity. In sum, these data showing that long-isoform-specific mutations are hypomorphic suggest that the UNC-5B short isoform has a role in axon guidance.

### *unc-5* hypomorphic mutations affect growth cone protrusions and polarity

During dorsal migration, VD growth cones display a robust lamellipodial growth cone body with filopodial protrusions biased to the dorsal region of the growth cone (KNOBEL *et al*. 1999; NORRIS AND LUNDQUIST 2011) (Figure 4A-D). The strong loss-of-function *unc-5(e53)* displayed growth cones that were excessively protrusive compared to wild-type, with larger growth cone area and longer filopodial protrusions (NORRIS AND LUNDQUIST 2011). Dorsal bias of filopodial protrusion was also lost in *unc-5(e53)* (NORRIS AND LUNDQUIST 2011). Strong loss-of-function *unc-5(e791)* showed a similar loss of filopodial dorsal polarity and excess protrusion (Figure 4 A-B, and F). The *unc-5(e152)* hypomorphic allele also displayed increased growth cone area and filopodial length, and loss of dorsal bias of filopodial protrusion (NORRIS AND LUNDQUIST 2011) (Figure 4), as did the *unc-5(ev480)* hypomorphic mutant (Figure 4).

**Figure 4.**
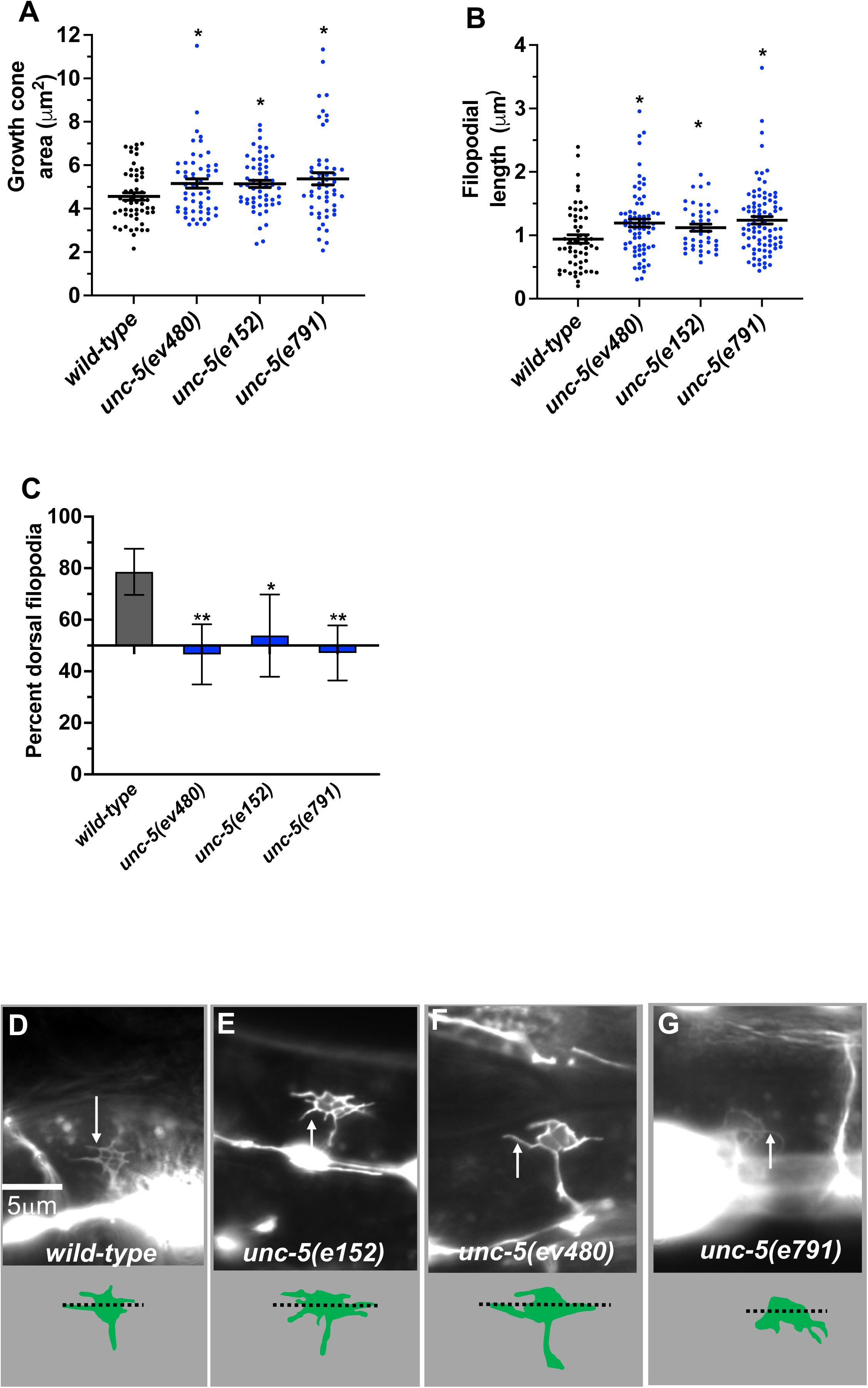
VD growth cone morphology in *unc-5* mutants. At least 50 growth cones were scored for each genotype. In the graphs, each point represents a measurement of a single growth cone or filopodium. (A, B) Quantification of VD growth cone area and filopodial length (see Materials and Methods). (A) Area of growth cone, in μm^2^. (B) Filopodial length, in μm. Error bars indicate standard error of the mean. Two-sided *t-* tests with unequal variance were used to determine the significance between wild-type and mutants. Single asterisks (*) indicate significance at *p* < 0.05. Double asterisks (**) indicate significance at *p* < 0.001. C) A graph showing the percent of dorsally directed filopodial protrusions in VD growth cones of different genotypes (see Materials and Methods). The X-axis is set at 50%, such that bars extending above the X-axis represents above 50%, and bars that extends below represents below 50%. In wild-type, a majority of filopodia (78%) extended from the dorsal half of the growth cone. Significance between wild-type and mutants was determined by Fisher’s exact test. Error bar represents 2x standard error of proportion. 50 growth cones were scored. Single asterisks (*) indicates the significant *p* < 0.05 double asterisks (**) indicate significant *p* < 0.001. (D-G) Fluorescence micrographs of wild-type and mutant VD growth cones expressing *Punc-25::gfp*. Arrows point to filopodial protrusions. Dorsal is up; anterior is left. The scale bar in (D) represents 5μm. Below each figure is a depiction of the growth cone with the dorsal and ventral regions separated by a dotted line.

Previous studies showed that F-actin in VD growth cones shows a dorsal bias of F-actin accumulation that is dependent upon UNC-5 (NORRIS AND LUNDQUIST 2011; NORRIS *et al*. 2014; GUJAR *et al*. 2018). Furthermore, UNC-5 was required to restrict the presence of microtubule + ends in growth cones, which have a pro-protrusive effect (GUJAR *et al*. 2018). These studies were conducted with strong loss-of-function alleles of *unc-5*.

Hypomorphic *unc-5(e152)* and *unc-5(ev480)* also displayed loss of dorsal F-actin accumulation (Figure 5A and D) and increased numbers of microtubule + ends in growth cones (Figure 5E and G). In sum, these results indicate that the UNC-5 long isoforms are required for growth cone dorsal polarity of filopodial protrusion and F-actin accumulation, to inhibit growth cone protrusion, and to restrict MT + end entry into the VD growth cone.

**Figure 5.**
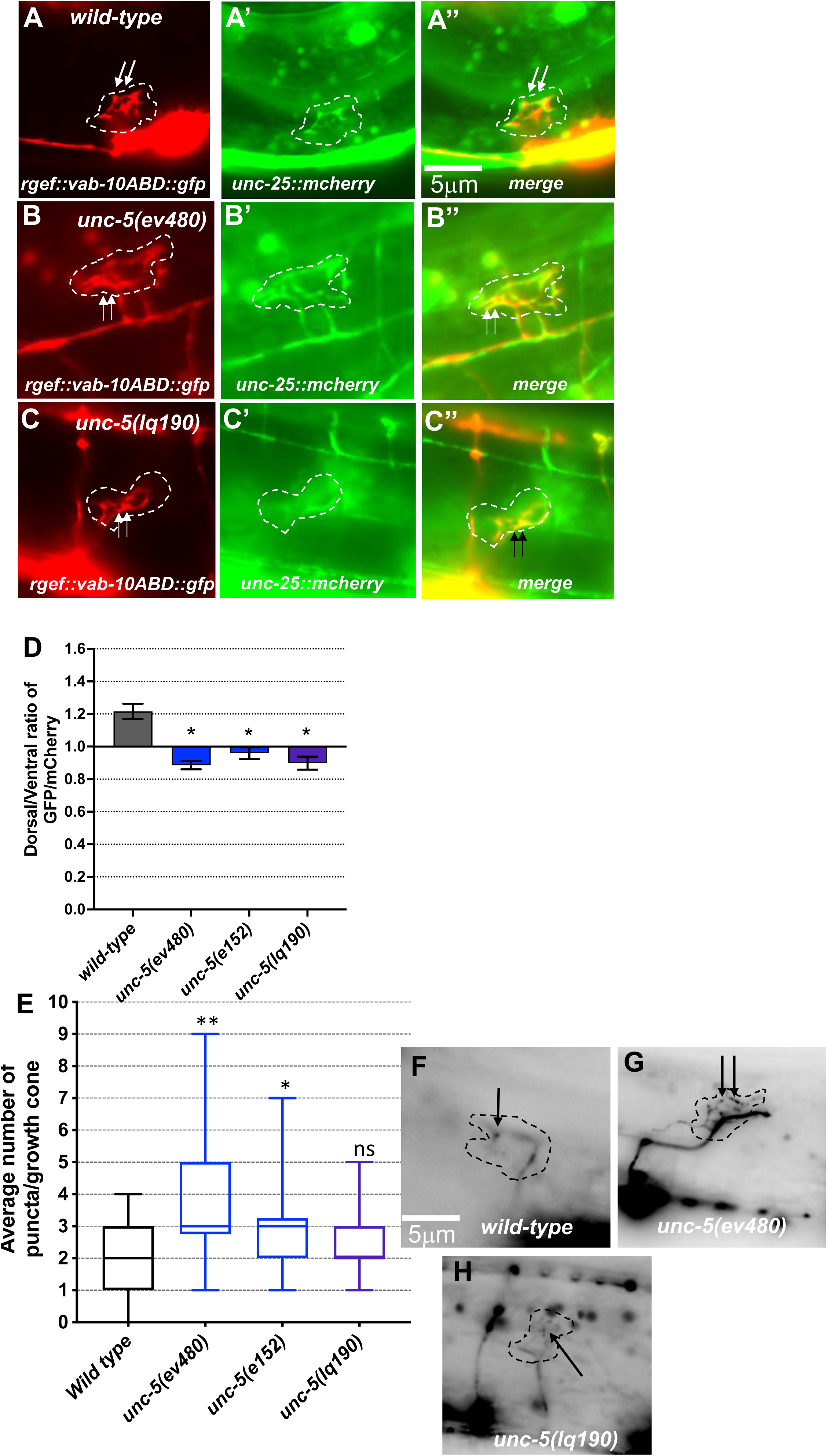
F-actin dorsal accumulation and microtubule + end distribution in VD growth cones. In all images, dorsal is up and anterior is left. Dashed lines indicate the growth cone perimeter. Scale bars represent 5 μm. (A) mCherry growth cone volume marker of a *wild-type* VD growth cone. (A’) VAB-10ABD::GFP accumulation at the dorsal edge of a wild-type VD growth cone (arrows) (see Materials and Methods). (A’’) Merge. (B, B’, B’’) As in A, for an *unc-5(ev480)* growth cone. (C, C’, C’’) As in A, for an *unc-5(lq190)* growth cone. (D) A graph of the dorsal/ventral ratio of GFP/mCherry from multiple growth cones (≥15) in *wild-type* and mutant animals expressing VAB-10ABD::GFP and mCherry (a volumetric marker) as described previously (NORRIS AND LUNDQUIST 2011) (see Materials and Methods). Error bars represent the standard error of the mean of the ratios from different growth cones. Growth cones were divided into dorsal and ventral halves, and the average intensity ratio of VAB-10ABD::GFP/mCherry was determined for each half and represented. Asterisks (*) indicate the significance of difference between wild-type and the mutant phenotype (\**p* < 0.05) (two-tailed *t*-test with unequal variance between the ratios of multiple growth cones of each genotype). (E) Box-and-whiskers plot of the number of EBP-2::GFP puncta in the growth cones of different genotypes (≥25 growth cones for each genotype). The boxes represent the upper and lower quartiles, and error bars represent the upper and lower extreme values. Asterisks (*) indicate the significant difference between wild-type and the mutant phenotype (\**p* < 0.05, \*\**p* < 0.001) determined by two-sided *t*-test with unequal variance. n.s., not significant. (F-H) Fluorescence micrographs of EBP-2 distribution in the VD growth cones of indicated genotypes. Dashed lines indicate the growth cone perimeter. Dorsal is up and anterior is left. Scale bar: 5μm.

### An *unc-5B* short isoform-specific mutation affects growth cone polarity and filopodial protrusion

A mutation predicted to affect only the *unc-5B* short isoform was constructed by CRISPR/ Cas9 genome editing. *unc-5B(lq190)* was produced by precise removal of intron 7, which is predicted to preclude the formation of the *unc-5B* short isoform by fusing together exons seven and eight (Figure 1A), leaving the long isoforms unaffected.

*unc-5B(lq190)* mutants displayed wild-type locomotion and weak but significant total axon defects (3%) including axon wandering (Table 1 and Figure 3K). *unc-5B(lq190)* did not significantly affect VD growth cone area (Figure 6A). However, filopodial length was reduced compared to wild-type (Figure 6B), and dorsal polarity of filopodial protrusion was abolished (Figure 6C and D). Consistent with the loss of dorsal filopodial polarity, F-actin dorsal accumulation was also lost in *unc-5B(lq190)* (Figure 5A and D). Microtubule + end accumulation was unaffected (Figure 5E and H).

**Figure 6.**
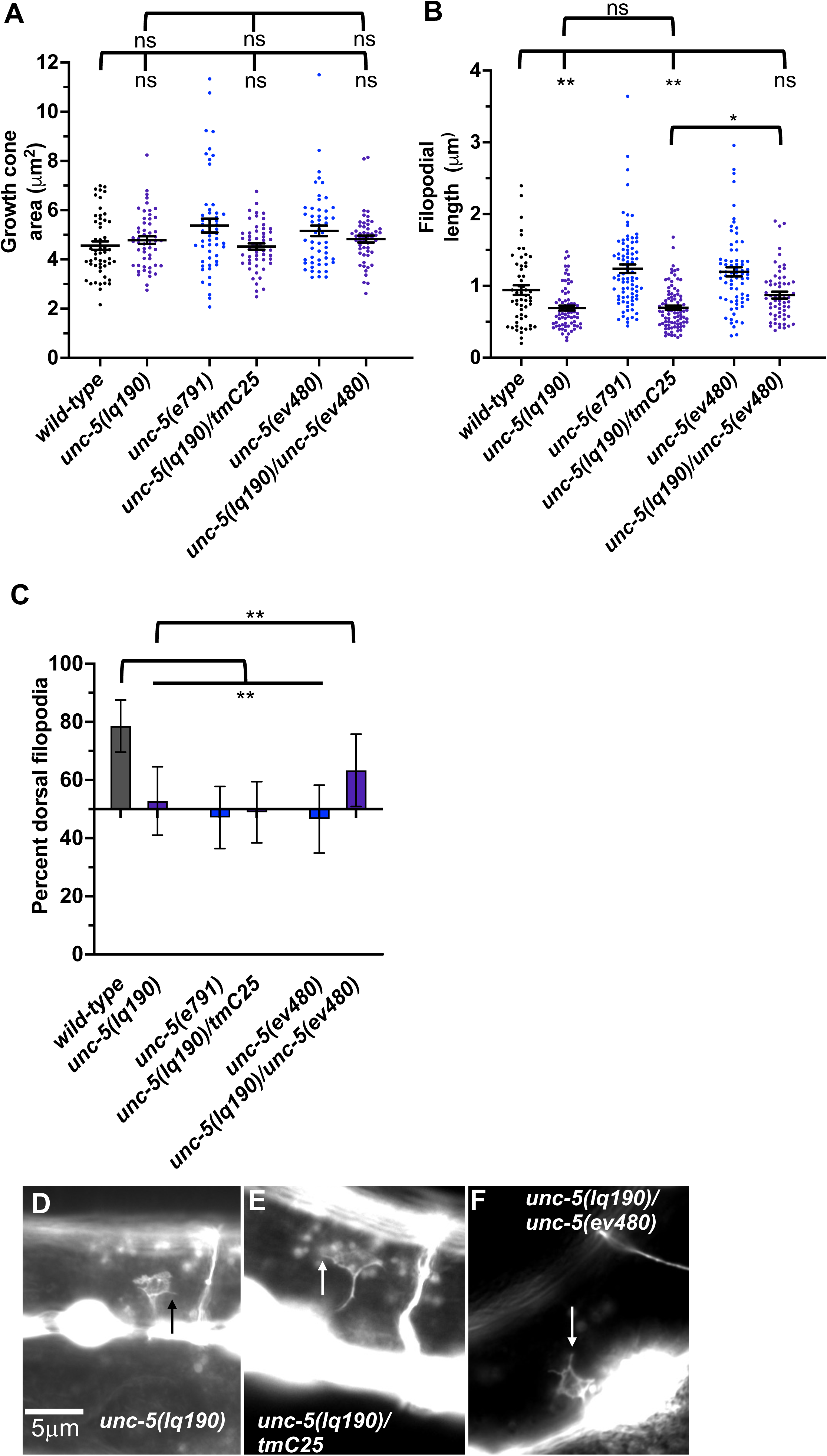
Growth cone protrusion and polarity in the short-isoform-specific *unc-5(lq190)* mutant. (A-C) Quantification of growth cone area, filopodial length, and dorsal polarity of filopodial protrusion as described in Figure 4. (D-F) Representative images of growth cones as described in Figure 4. Arrows point to filopodia.

To ensure that the phenotype observed in *unc-5B(lq190)* was due to the *unc-5* genome edit and not to a background mutation, we analyzed *trans-*heterozygotes of *unc-5B(lq190)* and the strong loss-of-function allele *unc-5(tmIs1241)* found on the balancer chromosome *tmC25* (DEJIMA *et al*. 2018)*. tmIs1241* is an insertion of a *myo-2::gfp* transgene in the *unc-5* exon seven shared by long and short isoforms (DEJIMA *et al*. 2018), and has a strong Unc phenotype and severe VD/DD axon guidance defects similar to *unc-5(e791)* (data not shown). Thus, *unc-5(tmIs1241)* a strong loss-of-function *unc-5* mutant. VD growth cones of *lq190/tmC25 trans-*heterozygotes resembled *unc-5B(lq190)* alone, with significantly reduced filopodial length compared to *wild-type* and loss of dorsal polarity of filopodial protrusion (Figure 6). This indicates that *unc-5(tm1241)* failed to complement *unc-5B(lq190)* as expected.

An *lq190 trans-*heterozygote with the hypomorphic *ev480* allele displayed filopodial length similar to *wild-type* and dorsally-polarized filopodial protrusion (Figure 6). Thus, *unc-5(ev480)* complemented *unc-5B(lq190)*, consistent with the idea that these isoforms encode distinct functions. In sum, these results indicate that *unc-5* long and *unc-5B* short encode genetically distinct and separable functions. *unc-5B* short isoform is required for dorsal polarity of filopodial protrusion, as well as robust filopodial protrusion, suggesting a pro-protrusive role.

### UNC-5A and UNC-5B both facilitate VD/DD axon guidance

The phenotypic analysis above suggests that the UNC-5 long isoforms and the UNC-5B short isoform are each involved in VD/DD axon guidance. An *unc-5::gfp* minigene is composed of the *unc-5* promoter driving the *unc-5A* cDNA fused to *gfp* (KILLEEN *et al*. 2002). This *unc-5A* minigene lacks introns and therefore is predicted to be unable to produce the *unc-5B* short isoform. Transgenic expression of *unc-5A* significantly rescued the Unc phenotype, VD/DD lateral midline crossing axon defects, and total VD/DD axon guidance defects of *unc-5* mutants, including *e791* and hypomorphic *e152* and *ev480* mutants (Table 1 and Figure 3G and I). *unc-5A* expression in a wild-type background resulted in a slightly uncoordinated phenotype with weak but significant VD/DD axon guidance defects (Table 1 and Figure 3E). This is possibly due to transgenic overexpression.

An *unc-5B* short isoform minigene was constructed by deleting exons eight and nine of the *unc-5A* minigene and replacing them the first 45 bases of intron 7 encoding the C-terminal 15 residues of UNC-5B short, followed by *gfp* (Figure 1B). This *unc-5B* minigene has the capacity to produce a molecule with the UNC-5B C-terminus, and not the longer isoforms. Transgenic *unc-5B* in a wild-type background also led to weak but significant VD/DD axon guidance defects and a slightly Unc phenotype (Table 1 and Figure 3F). *unc-5B* rescued the Unc, midline crossing, and total axon guidance defects of *e791, e152,* and *ev480*. However, rescue of total axon guidance defects was significantly weaker than *unc-5A* (Table 1 and Figure 3 F, H, and J). These data indicate that UNC-5A long and UNC-5B short can both facilitate VD/DD axon guidance.

### MYR::UNC-5B did not affect VD growth cone morphology

A constitutively-active UNC-5A transgene was produced by adding an N-terminal myristylation sequence to the cytoplasmic domain of UNC-5A (NORRIS *et al*. 2014). Expression of MYR::UNC-5A in the VD/DD neurons strongly inhibited VD growth cone area and filopodial protrusion length, consistent with a role of UNC-5A in inhibiting protrusion (NORRIS *et al*. 2014) (Figure 7A-B). MYR::UNC-5A expression caused no significant defects in polarity of protrusion (Figure 7C).

**Figure 7.**
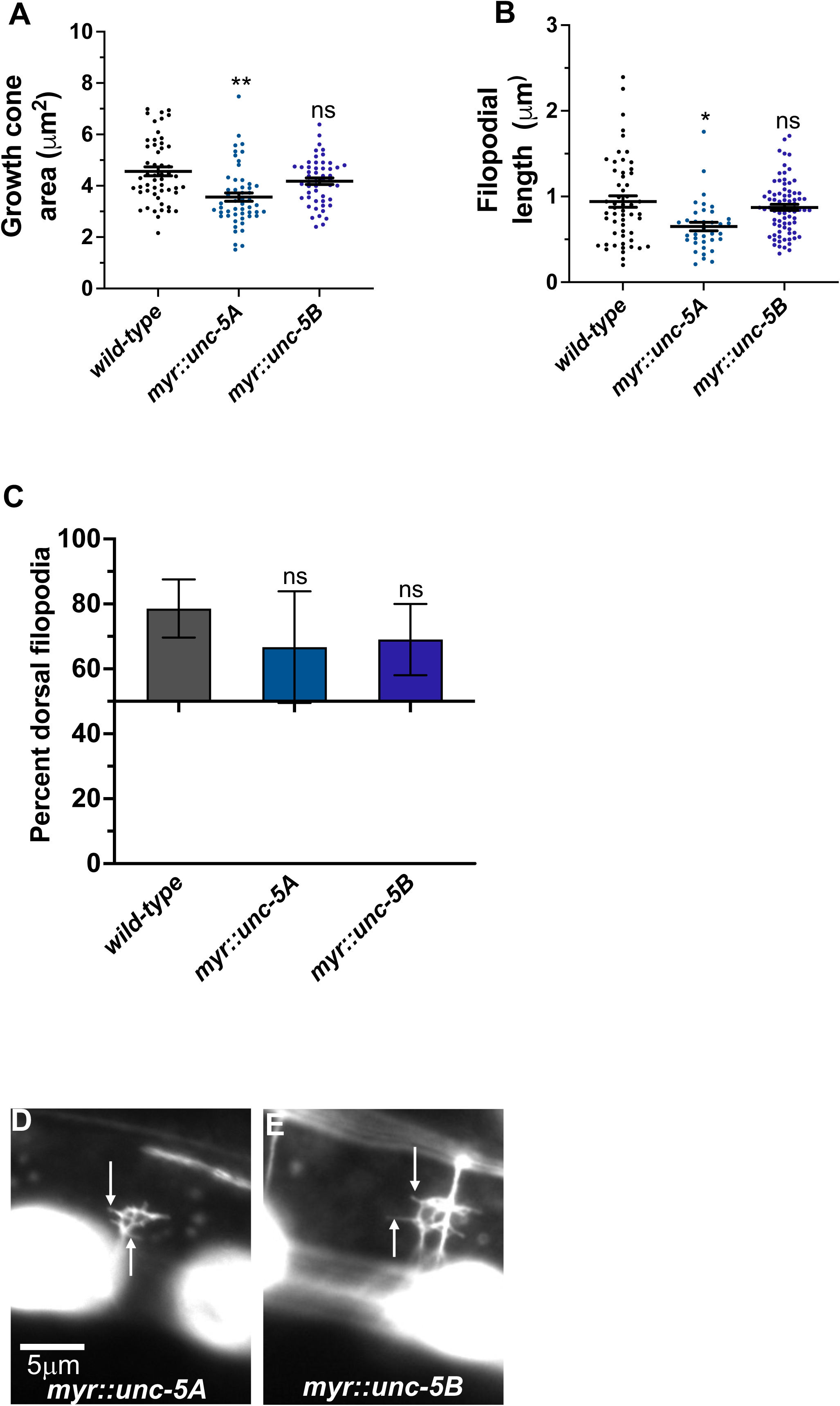
MYR::UNC-5B does not affect VD growth cone morphology. (A-C) Quantification of growth cone area, filopodial length, and dorsal polarity of filopodial protrusion as described in Figure 4. (D and E) Representative images of growth cones as described in Figure 4. Arrows point to filopodia.

A MYR::UNC-5B short isoform transgene was constructed with the 3’ end of *unc-5B* (Figure 2A). Expression of MYR::UNC-5B in VD/DD neurons caused no significant change in growth cone area, filopodial length, or polarity (Figure 7). This is consistent with *unc-5B(lq190)* short isoform having a distinct role from the long isoforms, which inhibit protrusion.

### *unc-5B* genetic interactions with *unc-6* and *unc-40*

The UNC-6/Netrin receptor UNC-40/DCC has a dual role in controlling growth cone protrusion. It acts as a heterodimer with UNC-5 to inhibit protrusion, and as a homodimer to stimulate protrusion (NORRIS AND LUNDQUIST 2011; NORRIS *et al*. 2014; GUJAR *et al*. 2018). *unc-40(n324)* null mutant displayed a slight but not significant reduction in growth cone area, and significantly shorter filopodial protrusions (Figure 8A-B and D). *unc-40(n324)* did not significantly affect dorsal polarity of filopodial protrusions (Figure 8C). This intermediate phenotype likely reflects the dual role of UNC-40 in stimulation and inhibition of VD growth cone protrusion.

**Figure 8.**
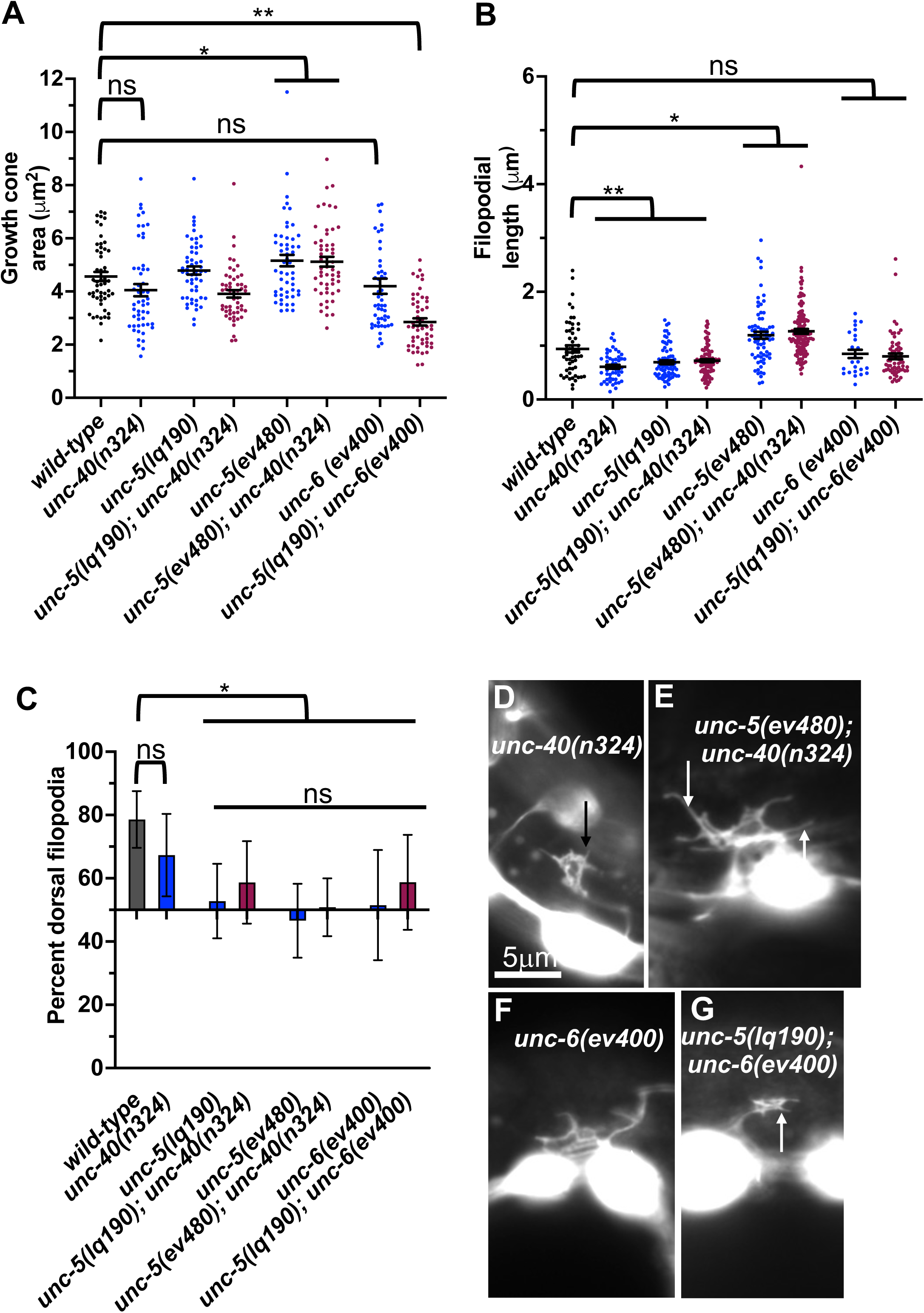
**VD growth cone morphology in *unc-5* hypomorphic mutants and interactions with *unc-40* and *unc-6.*** (A-C) Quantification of growth cone area, filopodial length, and dorsal polarity of filopodial protrusion as described in Figure 4. (D-G) Representative images of growth cones as described in Figure 4. Arrows point to filopodia.

Previous studies showed that UNC-40 function was required for the excess growth cone lamellipodial and filopodial protrusion seen in *unc-5* strong loss-of-function mutants that affect both long and short isoforms. Thus, in an *unc-5* mutant, the pro-protrusive activity of UNC-40 was overactive, resulting in excess protrusion (i.e. *unc-40* suppressed *unc-5*) (NORRIS AND LUNDQUIST 2011). *unc-5(lq190); unc-40(n324)* double mutant growth cones resembled the additive effect of each mutant alone: slight but not significant reduction in growth cone area; significantly reduced filopodial length, and loss of dorsal polarity of filopodial protrusion (Figure 8). This suggests that UNC-5B short isoform and UNC-40 might act in in the same pathway for filopodial length. However, *unc-5B(lq190)* synergistically increased *unc-40(n324)* VD/DD axon guidance defects (Table 1 and Figure 3 M), suggesting that they have might act in parallel in other aspects of growth cone morphology not assayed here.

*unc-6(ev400)* strong loss-of-function resulted in unpolarized growth cones with no significant effect on growth cone and filopodial protrusion (Figure 8). *unc-6(ev400); unc-5(lq190)* growth cones showed significantly reduced growth cone area compared to either single mutant alone. Filopodial length resembled the short filopodia of *unc-5(lq190)* alone. This suggests that *unc-6* and *unc-5B* short might act in parallel to promote growth cone protrusion and is consistent with a pro-protrusive role of *unc-5B* short isoform.

### *unc-40* did not suppress excess protrusion of an *unc-5* hypomorph

*unc-5* hypomorphic mutants that affect only the long isoforms also displayed excess growth cone protrusion (Figure 4). *unc-40(n324)* did not significantly reduce increased growth cone area or filopodial length in *unc-5(ev480)* hypomorphic mutant (Figure 8A-B and F). This suggests that the excess protrusion seen in the *unc-5(ev480)* hypomorph is not solely due to overactivity of UNC-40. Possibly, the UNC-5B short isoform has pro-protrusive activity that is increased in the absence of the long isoforms. This is consistent with *unc-5B(lq190)* short isoform mutants having reduced filopodial length.

Previous results showed that *unc-40* mutation suppressed the VD/DD axon guidance defects of *unc-5(e53)* strong loss-of-function and of *unc-5(e152)* hypomorphic mutants (NORRIS AND LUNDQUIST 2011). In contrast, the *unc-5(ev480)* hypomorph was not suppressed by *unc-40(n324)* and in fact showed synergistic enhancement for failure of lateral midline crossing and total axon guidance defects (Table 1 and Figure 3 N). This is consistent with an UNC-40-independent mechanism in *unc-5(ev480)* hypomorphic mutants driving excess protrusion, possibly involving UNC-5B.

### UNC-5A transgenic expression rescued VD growth cone defects in *unc-5* mutants

Transgenic expression of UNC-5A and UNC-5B each rescued the uncoordinated locomotion and VD/DD axon guidance defects on *unc-5(e791)* mutants (Table 1). UNC-5A expression in a *wild-type* background did not affect growth cone area or filopodial length (Figure 9A and B), but did significantly reduce dorsal polarity of filopodial protrusion, but not to the extent of *unc-5* mutants (Figure 9C). This might explain in part the axon guidance defects caused by *unc-5A* expression in a *wild-type* background (Table 1). UNC-5A expression significantly rescued excess growth cone area, filopodial length, and dorsal polarity of filopodial protrusion in *unc-5(e791)* mutants (Figure 9A-C).

**Figure 9.**
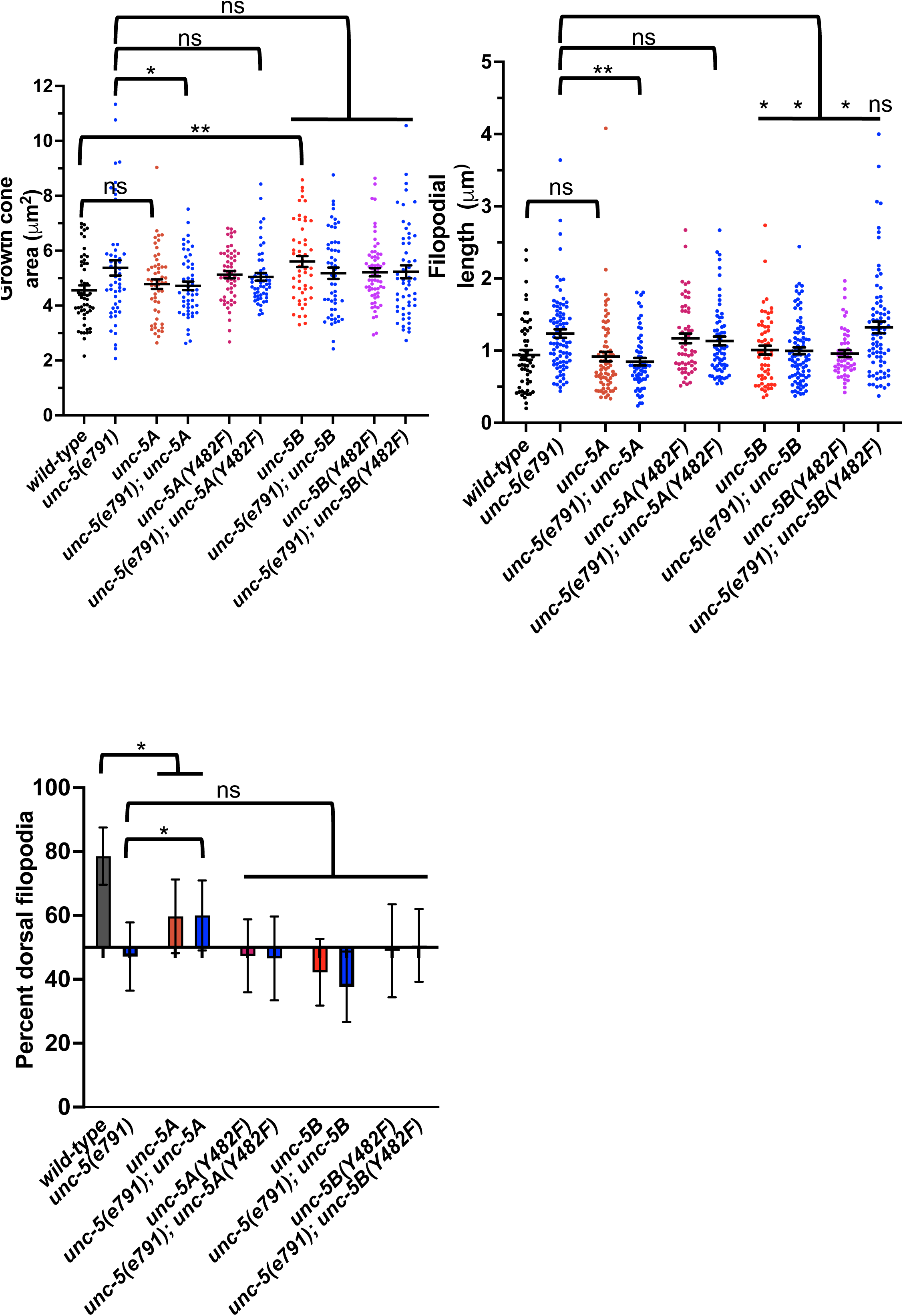
Effects of transgenic expression of UNC-5A and UNC-5B on VD growth cones. (A-C) Quantification of growth cone area, filopodial length, and dorsal polarity of filopodial protrusion as described in Figure 4. (D-F) Representative images of growth cones as described in Figure 4. Arrows point to filopodia.

### UNC-5B transgenic expression increased VD growth cone area and disrupted dorsal polarity of filopodial protrusion

UNC-5B expression in a *wild-type* background caused significantly increased growth cone area, but did not affect filopodial length (Figure 9A and B). Furthermore, UNC-5B expression abolished dorsal polarity of filopodial protrusion (Figure 9C). This might in part explain the VD/DD axon guidance defects caused by *unc-5B* expression (Table 1).

UNC-5B expression failed to significantly rescue growth cone area or dorsal polarity of filopodial protrusion in *unc-5(e791)* mutants (Figure 9A and B), but did significantly reduce filopodial length (Figure 9C). It is possible UNC-5B acts as a dominant-negative for UNC-5A activity. However, UNC-5B expression rescued uncoordinated locomotion and filopodial length of *unc-5* mutants, suggesting that it might have roles in the VD growth cone distinct from UNC-5A.

### Tyrosine 482 was required for UNC-5A function

Tyrosine 482 was previously shown to be required for the ability of an *unc-5* transgene to rescue *unc-5* mutants (KILLEEN *et al*. 2002). Tyrosine 482 was mutated to non-phosphorylable phenylalanine in *unc-5A* and *unc-5B* transgenes. Rescue of *unc-5(e791)* VD/DD axon guidance defects by *unc-5A(Y482F)* was significantly reduced compared to *unc-5A(+)* (Table 1).

In a wild-type background, *unc-5A(Y482F)* caused a significant increase in growth cone area and filopodial length similar to *unc-5(e791)* mutants (Figure 9A and B), and abolished dorsal polarity of filopodial protrusion (Figure 9C). Furthermore, it failed to significantly rescue growth cone area, filopodial length, and polarity of protrusion in *unc-5(e791)* mutants (Figure 9). These data indicate that Y482F is required for UNC-5A function, and that UNC-5A(Y482) might have a dominant negative effect on wild-type UNC-5A.

### Tyrosine 482 was required for some aspects of UNC-5B function

*unc-5B(Y482F)* failed to rescue VD/DD axon guidance defects in *unc-5* mutants, indicating that Y482 is important in UNC-5B function (Table 1). It also failed to rescue filopodial length defects of *unc-5* mutants (Figure 9B). UNC-5B(Y482F) expression alone still caused an increase in growth cone area and abolished dorsal polarity of filopodial protrusion (Figure 9A and C). This suggests that Y482 is required for some aspects of UNC-5B function (axon guidance and filopodial length) but not others (increased growth cone area and disruption of dorsal polarity of protrusion).

## Discussion

Studies reported here demonstrate a role for the novel and previously unreported UNC-5B short isoform in axon guidance and growth cone morphology. UNC-5B lacks most of the cytoplasmic domains of UNC-5 long isoforms, including the DEATH domain, the UPA domain, and most of the ZU5 domain (Figures 1 and 2). Both long and short isoforms are required for dorsal polarity of VD growth cone filopodial protrusion. However, in contrast to the long isoforms, which inhibit VD growth cone protrusion, UNC-5B short might be required for robust VD growth cone lamellipodial and filopodial protrusion.

### Mutations specifically affecting the *unc-5* long isoforms are hypomorphic

*unc-5* mutations that affect all *unc-5* caused severely uncoordinated locomotion and severe VD/DD axon guidance defects (*unc-5* strong loss-of-function). Very few axons were observed emanating from the ventral nerve cord, and those that did failed to extend dorsally past the lateral midline (Figure 3 and Table 1). *unc-5* mutations that affect only the long isoforms were less severely uncoordinated than *unc-5* strong loss-of-function mutants, and displayed less-severe VD/DD axon guidance defects (*unc-5* hypomorphs) (Table 1 and Figure 1). However, VD growth cone morphology of *unc-5* hypomorphs was not significantly different from that of *unc-5* strong loss-of-function. They were unpolarized with larger growth cones and longer filopodial protrusions, and displayed unpolarized F-actin and excess microtubule + ends. It is likely that subtle differences in VD growth cone morphology between strong and hypomorphic *unc-5* mutants were not detected in these assays. In any case, the weaker uncoordination and weaker axon guidance defects of *unc-5* hypomorphs imply that the *unc-5B* short isoform has a role in VD/DD axon guidance.

### UNC-5B short is required for dorsal polarity of VD growth cone filopodial protrusion and for robust filopodial protrusion

A mutation precited to specifically affect the *unc-5B* short isoform was generated by CRISPR/Cas9 genome editing. The *unc-5B* 3’ end results from failure to splice intron 7 from the *unc-5* transcript, resulting in a stop codon after 16 additional amino acid coding codons (Figure 1). RNA-seq revealed *unc-5B* reads in intron 7 (Figure 1). *unc-5B(lq190)* was a precise removal of intron 7, resulting in the fusion of exons 7 and 8, leaving *unc-5* long isoform coding potential unchanged. *unc-5b(lq190)* mutants were only slightly uncoordinated with weak VD/DD axon guidance defects (Table 1 and Figure 3). However, *unc-5B* VD growth cones were unpolarized and displayed significantly shorter filopodial protrusions compared to wild-type (Figure 6). *unc-5B(lq190)* failed to complement an *unc-5* strong loss-of-function mutation for these phenotypes, but did complement an *unc-5* hypomorphic long isoform-specific mutant (Figure 6). Thus, the functions of *unc-5* long and short isoforms are genetically distinct and separable. *unc-5B(lq190)* resulted in unpolarized F-actin, but did not affect microtubule + end abundance, consistent with the growth cone phenotypes of loss of polarity but no effect on growth cone area.

These data suggest that UNC-5B short has a pro-protrusive role, in contrast to the inhibitory role of UNC-5 long. Consistent with this idea, *unc-5B(lq190)* double mutants with *unc-6* had severely reduced VD growth cone area compared to wild-type and single mutants alone, and transgenic expression of *unc-5B* resulted in significantly increased growth cone area compared to wild type (Figure 9).

### Transgenic expression of UNC-5B short partially rescues *unc-5* mutants

Transgenic expression of *unc-5A* long caused slightly uncoordinated locomotion and weak VD/DD axon guidance defects (Table 1 and Figure 3). *unc-5A* long expression did not significantly alter growth cone area or filopodial length, but did significantly perturb dorsal polarity of protrusion, but not to the extent of *unc-5* strong loss-of-function mutants (Figure 9). *unc-5A* long transgenic expression did significantly rescue the uncoordinated locomotion and VD/D axon guidance defects of *unc-5* strong loss-of-function mutants, as well as VD growth cone area and filopodial length increase (Table 1, Figure 3, and Figure 9). Thus, while *unc-5A* long transgenic expression caused defects alone, it did rescue *unc-5* loss-of-function.

Transgenic *unc-5B* short expression also caused weak uncoordinated locomotion and VD/DD axon guidance defects, as well as loss of dorsal polarity of protrusion and increased growth cone area (Table 1, Figure 3, and Figure 9). *unc-5B* short transgenic expression rescued uncoordinated locomotion and VD/DD axon guidance defects of *unc-5* mutants, but did not rescue polarity or growth cone area defects of *unc-5* strong loss-of-function. This indicates that more subtle effects of *unc-5B* short expression on the growth cone are not being resolved in these studies. *unc-5B* short expression did rescue excess filopodial length of *unc-5* strong loss-of-function (Figure 9), suggesting that in some contexts UNC-5B short might also inhibit filopodial protrusion. These results are consistent with previous studies showing that *unc-5* transgenes lacking cytoplasmic domains still retained function in axon guidance (KILLEEN *et al*. 2002).

These data indicate that both *unc-5A* long and *unc-5B* short can compensate for some aspects of *unc-5* loss-of function in axon guidance and that UNC-5B short has a novel role in axon guidance and growth cone morphology.

### Tyrosine 482 is required for UNC-5A long and UNC-5B short isoform function

Previous studies showed that tyrosine 482 (Y482) is phosphorylated and that it is required for the function of UNC-5 in axon guidance (KILLEEN *et al*. 2002). Studies here confirm that Y482 is required for the rescuing effects of the *unc-5A* long transgene. *unc-5(Y482F)* does not rescue the uncoordinated locomotion or VD/DD axon defects of *unc-5* mutants as well as the wild-type transgene (Table 1). Furthermore, it fails to rescue the increased growth cone area, excessively long growth cone filopodia, and loss of dorsal polarity of protrusion of *unc-5* mutants (Figure 9). Thus, Y482 is required for the function of UNC-5A long isoform.

Y482 was required for rescue of axon guidance and increased filopodial length of *unc-5* mutants. However, *unc-5B(Y482F)* expression alone still resulted in increased growth cone area and loss of filopodial protrusion. Thus, in UNC-5B short, Y482 is required for some functions but not others. While it is unclear which tyrosine kinase might phosphorylate Y482, the SH2 domain of SRC-1 binds to phosphorylated Y482, SRC-1 is required for robust UNC-5 tyrosine phosphorylation, and SRC-1 functions with UNC-5 in gonadal distal tip cell migration and axon guidance (LEE *et al*. 2005). Indeed, replacing the entire UNC-5 cytoplasmic region with SRC-1 in a fusion protein rescued *unc-5* mutants (LEE *et al*. 2005), suggesting that recruitment of SRC-1 is the role of the cytoplasmic domain of UNC-5. However, we show here that the UNC-5B short isoform lacking most of the cytoplasmic region of UNC-5 partially rescues *unc-5* mutants, so SRC-1 might not be required.

### UNC-5B short is a novel function of the *unc-5* locus in UNC-6/Netrin signaling in axon guidance

These results indicate that the UNC-5B short isoform is required for VD growth cone dorsal polarity of filopodial protrusion and for robust growth cone lamellipodial and filopodial protrusion. This is in contrast to the UNC-5A long isoform, which inhibits growth cone protrusion. Double mutants of *unc-5B(lq190); unc-40* displayed an additive phenotype, and no significant genetic interaction. This suggests that UNC-5B short might act independently of UNC-40, or in the same pathway. UNC-5B short is lacking the cytoplasmic UPA/DB domain that mediates physical interaction with the UNC-40 P1 cytoplasmic domain. This suggests that UNC-5B short might act independently of UNC-40.

*unc-5B(lq190); unc-6* double mutants displayed growth cones that were smaller than wild-type and either single mutant alone. This suggests that UNC-5B short might act in parallel to UNC-6 to promote growth cone lamellipodial protrusion. In the Polarity/Protrusion model, UNC-6 acts through UNC-40 to stimulate growth cone protrusion. However, this effect is independent of UNC-40 as *unc-5B(lq190); unc-40* double mutants did not display small growth cones similar to *unc-5B(lq190); unc-6* mutants. This suggests that an unidentified ligand might interact with UNC-5B short to stimulate protrusion in parallel to UNC-6, and that an unidentified receptor for UNC-6 might act in parallel to UNC-5B short (Figure 10). The identities of this potential and ligand are unknown, but could other cues and receptors that guide axons in the dorsal ventral axis and interact with UNC-6/netrin signaling, such as Slit/Robo (TANG AND WADSWORTH 2014; XU *et al*. 2015), Draxin (GAO *et al*. 2015), or dsCAM (PUROHIT *et al*. 2012).

**Figure 10.**
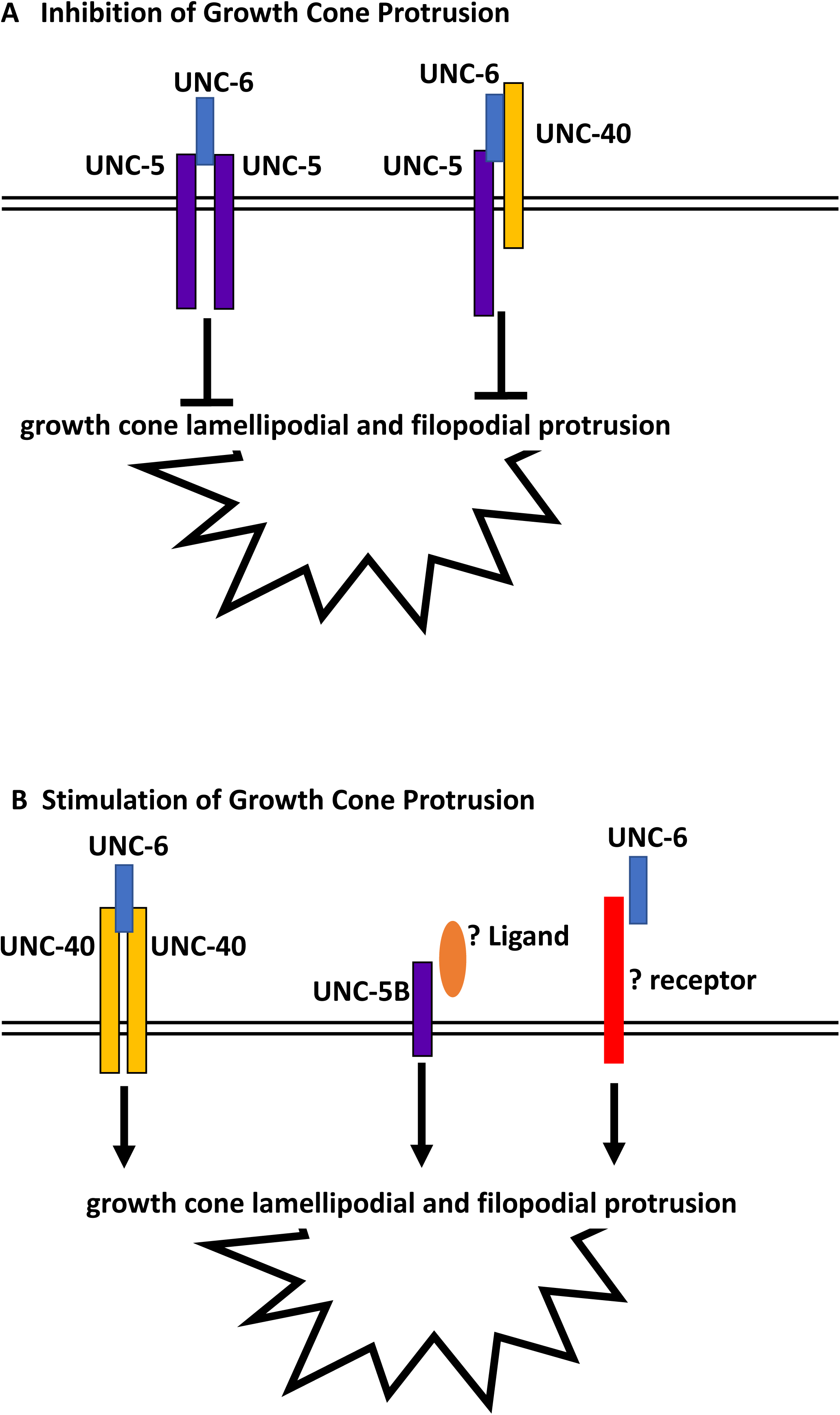
Regulation of growth cone protrusion by UNC-6/Netrin and its receptors. A) UNC-6/netrin inhibits VD growth cone protrusion through UNC-5 homodimers and UNC-5:UNC-40 heterodimers. B) UNC-6/Netrin stimulates growth cone protrusion via UNC-40 homodimers. Work described here shows that UNC-6/Netrin acts in parallel to the novel UNC-5B short isoform to stimulate growth cone protrusion in an UNC-40-independent manner.

In sum, this work describes a novel role of a short UNC-5 isoform lacking most cytoplasmic domains in axon guidance and growth cone morphology. Previous studies showed that UNC-5 homodimers and UNC-5:UNC-40 heterodimers inhibit VD growth cone protrusion, and that UNC-40 homodimers stimulate growth cone protrusion in response to UNC-6 (Figure 10) (NORRIS AND LUNDQUIST 2011; NORRIS *et al*. 2014; Gujar *et al*. 2018; Gujar *et al*. 2019; Mahadik and Lundquist 2022; Mahadik and LUNDQUIST 2023). These results show that the UNC-5B short isoform is pro-protrusive, acts independently of UNC-40, and in parallel to UNC-6.

## Material and Methods

### Genetic Methods

Experiments were performed at 20^0^C using standard *C. elegans* techniques (BRENNER 1974). The presence of mutations was confirmed by phenotype and sequencing. The following integrated transgenes were produced, with unknown chromosomal location: *lqIs283 [Punc-5::unc-5A]*, *lqIs308 [Punc-5::unc-5B], lqIs360 [Punc-5::unc-5A(Y482F)], lqIs387 [Punc-5::unc-5A(Y482F)], lqIs326 [Punc-5::myr::unc-5A], lqIs315 [Punc-5::myr::unc-5B]*.

### Transgene construction

The PUNC5GFP transgene (KILLEEN *et al*. 2002) was used as a basis to construct the transgenes described in this work. The *Punc-5::unc-5B* short isoform transgene was constructed by removing the coding sequence of exons eight and nine, and replacing with the first 45 bases of intron 7 (Figure 1) fused in-frame to *gfp*. The tyrosine 482 mutation to phenylalanine was generated by site-directed mutagenesis. Sequences of the transgenes are in Supplemental File 1.

### RNA-seq and analysis

RNA-seq was conducted using total RNA isolated from three independent isolates of mixed-stage animals of the strain LE6194 (*wrdSi23 I; juIs76 II*). (SSM1, SSM2, and SSM3) as previously described (TAMAYO *et al*. 2013). LE6194 is wild-type for the *unc-5* gene. Stranded poly-A RNA-seq libraries were made using the NEBNext® Ultra™ II Directional RNA Library Prep Kit for Illumina. Paired-end 150 cycle sequencing on a Nextseq550 was utilized. Reads were aligned to the *C. elegans* genome using HISAT2 with default settings (version 2.1.0) (KIM *et al*. 2019). Resulting BAM files were analyzed in the Integrated Genome Viewer (ROBINSON *et al*. 2011; THORVALDSDOTTIR *et al*. 2012) (Figure 1C). Sequences are available in the Sequence Read Archive, BioProject number PRJNA847250.

### Cas9 genome editing to generate *unc-5(lq190)*

*unc-5(lq190)* was generated by CRISPR/Cas9 genome editing which precisely deleted the entire intron 7 of *unc-5A,* between exon 7 and 8 (Figure 1 and Supplemental File 2). This removes the truncated 3’end of UNC-5B, which resides in intron 7, and leaves the coding potential of *unc-5* long isoforms unchanged. Synthetic guide RNAs were directed at the 5’ and 3’ ends of intron 7 (Supplemental File 2). A mix of sgRNAs, a single stranded oligonucleotide repair template with recoded sgRNA regions (Supplemental File 2), and Cas9 enzyme was injected into the gonads of N2 animals, along with the *dpy-10(cn64)* co-CRISPR reagents (EL MOURIDI *et al*. 2017). Deletion of *unc-5* intron 7 was confirmed by PCR and sequencing and outcrossed to N2 three times. Genome editing reagents were produced by InVivo Biosystems (Eugene, OR, USA):

sgRNA1: 5’ CTCTCCCCAGTCATCGTCAT 3’

sgRNA2: 5’ TAACAGATCAACCTCACCTC 3’

ssODN repair template (sequence of the *lq190* mutation): 5’GAAAATGCTTTATCTGGCTGTTTCGGACACCTTAA<u>CCGACCAGCCACATCTTAAGCCAATTG</u> <u>AGTCTGCCCTTTCTCCTGTTATTGT</u>CATCGGACAATGTGATGTTTCAATGTCAGCTCATG3’

(recoded exon sequence is underlined).

### Statistics

Proportional data were analyzed using Fisher’s Exact test. Continuous data were analyzed with Student’s *t*-test. Sample sizes used each gave an estimated statistical power of > 80% (a β false negative value of <0.2), with an LZ false positive value of 0.05.

### Imaging and quantification of axon guidance defects

VD/DD axons were visualized with *Punc-25*::*gfp* transgene, *juIs76* (JIN *et al*. 1999), which is expressed in 13 VD and 6DD GABAergic motor neurons. Axon guidance defects were scored as previously described (MAHADIK AND LUNDQUIST 2022). An average of 16 of the 19 commissures of VD/DD axons are distinguishable in wild-type, due to fasciculation and inability to distinguish individual axons. A total of 100 animals were scored (1600 total commissural processes). At a false positive *a* value of 0.05 with observed variances of 5% or greater, scoring 1600 axons gives an estimated statistical power of ≥ 80%. Total axon guidance defects were calculated by counting all the axons which failed to reach the dorsal nerve cord, wandered at an angle of 45^0^ or greater, crossed over other processes, and displayed ectopic branching. Significance difference between two genotypes was determined by using Fisher’s exact test.

### Growth cone imaging and quantification

Growth cones of VD axons were imaged as previously described (NORRIS AND Lundquist 2011; Norris *et al*. 2014; Gujar *et al*. 2017; Gujar *et al*. 2018; Gujar *et al*. 2019; MAHADIK AND LUNDQUIST 2022). Late L1/early L2 larval animals placed on a 2% agarose pad with 5mM sodium azide in M9 buffer. At n = 50, at a false positive a value of 0.05, statistical power of β (false negative) is much lower than 0.2.

Growth cones were imaged with Qimaging Retiga EXi camera on a Leica DM5500 microscope at 100x magnification. Projections less than 0.5um in width were scored as filopodia. Growth cone area and filopodial length were quantified using ImageJ software as previously described (NORRIS AND LUNDQUIST 2011; NORRIS *et al*. 2014; GUJAR *et al*. 2017; GUJAR *et al*. 2018; GUJAR *et al*. 2019; MAHADIK AND LUNDQUIST 2022). Significance of difference between two genotypes was determined by two-sided *t*-test with unequal variance.

Polarity of growth filopodial protrusions was determined as previously described (NORRIS AND LUNDQUIST 2011; NORRIS *et al*. 2014; GUJAR *et al*. 2017; GUJAR *et al*. 2018; GUJAR *et al*. 2019; MAHADIK AND LUNDQUIST 2022). Growth cone images were divided into dorsal and ventral halves with respect to the ventral nerve cord. The number of filopodia in each was counted. Proportion of dorsal filipodia was determined by the number of dorsal filopodia divided by total number of filopodia. Significance of difference between two genotypes was determined by Fisher’s exact test.

### VAB-10ABD::GFP imaging

Growth cone F-actin was analyzed as previously described (GUJAR *et al*. 2018), using the F-actin binding domain of VAB-10/spectraplakin fused to GFP (BOSHER *et al*. 2003; PATEL *et al*. 2008). A soluble mCherry volume marker was included in the strain. Growth cones images were captured as described above. ImageJ was used image analysis to determine asymmetric VAB-10ABD::GFP localization. The pixel intensities for VAB-10ABD::GFP were normalized to the volumetric mCherry fluorescence in five line scans from the dorsal half and the ventral half of each growth cone. This normalized ratio was determined for multiple growth cones, and the average and standard error for multiple growth cones was determined. Statistical comparisons between genotypes were done using a two-tailed *t-*test with unequal variance on these average normalized ratios of multiple growth cones of each genotype.

### EBP-2::GFP imaging

Growth cone EBP-2::GFP puncta (SRAYKO *et al*. 2005; KOZLOWSKI *et al*. 2007; YAN *et al*. 2013) was analyzed as described previously (GUJAR *et al*. 2018). Growth cone perimeter and filopodia were defined, and the EBP-2::GFP puncta in the growth cone were counted. Significance of difference was determined by a two-sided *t*-test with unequal variance.

### Data availability

The authors affirm that all data necessary for confirming the conclusions of the article are present within the article, figures, and tables. RNA seq FASTQ files are available in the Sequence Read Archive, BioProject number PRJNA847250.

## Acknowledgments

The authors thank E. Struckhoff, Z. Grant, and C. McKimens for technical assistance. Some strains were provided by the CGC, which is funded by NIH Office of Research Infrastructure Programs (P40OD010440). Sequencing was conducted at the KU Genome Sequencing Core supported by the National Institute of General Medical Sciences Center for Molecular Analysis of Disease Pathways (P30GM145499). The Kansas Infrastructure Network of Biomedical Research Excellence (P20GM103418) provided computational support. This work was supported by National Institutes of Neurological Disorders and Stroke grants R03NS114554 and R01NS115467 to E.A.L.

## References

1. Ackerman, S. L., L. P. Kozak, S. A. Przyborski, L. A. Rund, B. B. Boyer et al., 1997 The mouse rostral cerebellar malformation gene encodes an UNC-5-like protein. Nature 386: 838–842.

2. Bosher, J. M., B. S. Hahn, R. Legouis, S. Sookhareea, R. M. Weimer et al., 2003 The Caenorhabditis elegans vab-10 spectraplakin isoforms protect the epidermis against internal and external forces. J Cell Biol 161: 757–768.

3. Boyer, N. P., and S. L. Gupton, 2018 Revisiting Netrin-1: One Who Guides (Axons). Front Cell Neurosci 12: 221.

4. Brenner, S., 1974 The genetics of Caenorhabditis elegans. Genetics 77: 71–94.

5. Chan, S. S., H. Zheng, M. W. Su, R. Wilk, M. T. Killeen et al., 1996 UNC-40, a C. elegans homolog of DCC (Deleted in Colorectal Cancer), is required in motile cells responding to UNC-6 netrin cues. Cell 87: 187–195.

6. Dejima, K., S. Hori, S. Iwata, Y. Suehiro, S. Yoshina et al., 2018 An Aneuploidy-Free and Structurally Defined Balancer Chromosome Toolkit for Caenorhabditis elegans. Cell Rep 22: 232–241.

7. Dominici, C., J. A. Moreno-Bravo, S. R. Puiggros, Q. Rappeneau, N. Rama et al., 2017 Floor-plate-derived netrin-1 is dispensable for commissural axon guidance. Nature 545: 350–354.

8. El Mouridi, S., C. Lecroisey, P. Tardy, M. Mercier, A. Leclercq-Blondel, et al., 2017 Reliable CRISPR/Cas9 Genome Engineering in Caenorhabditis elegans Using a Single Efficient sgRNA and an Easily Recognizable Phenotype. G3 (Bethesda) 7: 1429–1437.

9. Gallo, G., and P. C. Letourneau, 1999 Axon guidance: A balance of signals sets axons on the right track. Curr Biol 9: R490–492.

10. Gao, X., U. Metzger, P. Panza, P. Mahalwar, S. Alsheimer et al., 2015 A Floor-Plate Extracellular Protein-Protein Interaction Screen Identifies Draxin as a Secreted Netrin-1 Antagonist. Cell Rep 12: 694–708.

11. Gujar, M. R., A. M. Stricker and E. A. Lundquist, 2017 Flavin monooxygenases regulate Caenorhabditis elegans axon guidance and growth cone protrusion with UNC-6/Netrin signaling and Rac GTPases. PLoS Genet 13: e1006998.

12. Gujar, M. R., A. M. Stricker and E. A. Lundquist, 2019 RHO-1 and the Rho GEF RHGF-1 interact with UNC-6/Netrin signaling to regulate growth cone protrusion and microtubule organization in Caenorhabditis elegans. PLoS Genet 15: e1007960.

13. Gujar, M. R., L. Sundararajan, A. Stricker and E. A. Lundquist, 2018 Control of Growth Cone Polarity, Microtubule Accumulation, and Protrusion by UNC-6/Netrin and Its Receptors in Caenorhabditis elegans. Genetics 210: 235–255.

14. Hedgecock, E. M., J. G. Culotti and D. H. Hall, 1990 The unc-5, unc-6, and unc-40 genes guide circumferential migrations of pioneer axons and mesodermal cells on the epidermis in C. elegans. Neuron 4: 61–85.

15. Ishii, N., W. G. Wadsworth, B. D. Stern, J. G. Culotti and E. M. Hedgecock, 1992 UNC-6, a laminin-related protein, guides cell and pioneer axon migrations in C. elegans. Neuron 9: 873–881.

16. Jin, Y., E. Jorgensen, E. Hartwieg and H. R. Horvitz, 1999 The Caenorhabditis elegans gene unc-25 encodes glutamic acid decarboxylase and is required for synaptic transmission but not synaptic development. J Neurosci 19: 539–548.

17. Killeen, M., J. Tong, A. Krizus, R. Steven, I. Scott et al., 2002 UNC-5 function requires phosphorylation of cytoplasmic tyrosine 482, but its UNC-40-independent functions also require a region between the ZU-5 and death domains. Dev Biol 251: 348–366.

18. Kim, D., J. M. Paggi, C. Park, C. Bennett and S. L. Salzberg, 2019 Graph-based genome alignment and genotyping with HISAT2 and HISAT-genotype. Nat Biotechnol 37: 907–915.

19. Knobel, K. M., E. M. Jorgensen and M. J. Bastiani, 1999 Growth cones stall and collapse during axon outgrowth in Caenorhabditis elegans. Development 126: 4489–4498.

20. Kozlowski, C., M. Srayko and F. Nedelec, 2007 Cortical microtubule contacts position the spindle in C. elegans embryos. Cell 129: 499–510.

21. Lee, J., W. Li and K. L. Guan, 2005 SRC-1 mediates UNC-5 signaling in Caenorhabditis elegans. Mol Cell Biol 25: 6485–6495.

22. Leonardo, E. D., L. Hinck, M. Masu, K. Keino-Masu, S. L. Ackerman et al., 1997 Vertebrate homologues of C. elegans UNC-5 are candidate netrin receptors. Nature 386: 833–838.

23. Leung-Hagesteijn, C., Spence, A.M., Stern, B.D., Zhou, Y., Su, M.-W., Hedgecock, E.M., Culotti, J.G., 1992 UNC-5, a transmembrane protein with immunoglobulin and thrombospondin type I domains, guides cell and pioneer axon migrations. Cell 71: 289–299.

24. Mahadik, S. S., and E. A. Lundquist, 2022 The PH/MyTH4/FERM molecule MAX-1 inhibits UNC-5 activity in the regulation of VD growth cone protrusion in Caenorhabditis elegans. Genetics 221.

25. Mahadik, S. S., and E. A. Lundquist, 2023 TOM-1/tomosyn acts with the UNC-6/netrin receptor UNC-5 to inhibit growth cone protrusion in Caenorhabditis elegans. Development 150.

26. Morales, D., 2018 A new model for netrin1 in commissural axon guidance. J Neurosci Res 96: 247–252.

27. Norris, A. D., and E. A. Lundquist, 2011 UNC-6/netrin and its receptors UNC-5 and UNC-40/DCC modulate growth cone protrusion in vivo in C. elegans. Development 138: 4433–4442.

28. Norris, A. D., L. Sundararajan, D. E. Morgan, Z. J. Roberts and E. A. Lundquist, 2014 The UNC-6/Netrin receptors UNC-40/DCC and UNC-5 inhibit growth cone filopodial protrusion via UNC-73/Trio, Rac-like GTPases and UNC-33/CRMP. Development 141: 4395–4405.

29. Patel, F. B., Y. Y. Bernadskaya, E. Chen, A. Jobanputra, Z. Pooladi et al., 2008 The WAVE/SCAR complex promotes polarized cell movements and actin enrichment in epithelia during C. elegans embryogenesis. Dev Biol 324: 297–309.

30. Przyborski, S. A., B. B. Knowles and S. L. Ackerman, 1998 Embryonic phenotype of Unc5h3 mutant mice suggests chemorepulsion during the formation of the rostral cerebellar boundary. Development 125: 41–50.

31. Purohit, A. A., W. Li, C. Qu, T. Dwyer, Q. Shao et al., 2012 Down syndrome cell adhesion molecule (DSCAM) associates with uncoordinated-5C (UNC5C) in netrin-1-mediated growth cone collapse. J Biol Chem 287: 27126–27138.

32. Robinson, J. T., H. Thorvaldsdottir, W. Winckler, M. Guttman, E. S. Lander et al., 2011 Integrative genomics viewer. Nat Biotechnol 29: 24–26.

33. Srayko, M., A. Kaya, J. Stamford and A. A. Hyman, 2005 Identification and characterization of factors required for microtubule growth and nucleation in the early C. elegans embryo. Dev Cell 9: 223–236.

34. Tamayo, J. V., M. Gujar, S. J. Macdonald and E. A. Lundquist, 2013 Functional transcriptomic analysis of the role of MAB-5/Hox in Q neuroblast migration in Caenorhabditis elegans. BMC Genomics 14: 304.

35. Tang, X., and W. G. Wadsworth, 2014 SAX-3 (Robo) and UNC-40 (DCC) regulate a directional bias for axon guidance in response to multiple extracellular cues. PLoS One 9: e110031.

36. Thorvaldsdottir, H., J. T. Robinson and J. P. Mesirov, 2012 Integrative Genomics Viewer (IGV): high-performance genomics data visualization and exploration. Brief Bioinform.

37. Varadarajan, S. G., and S. J. Butler, 2017 Netrin1 establishes multiple boundaries for axon growth in the developing spinal cord. Dev Biol 430: 177–187.

38. Wadsworth, W. G., H. Bhatt and E. M. Hedgecock, 1996 Neuroglia and pioneer neurons express UNC-6 to provide global and local netrin cues for guiding migrations in C. elegans. Neuron 16: 35–46.

39. Xu, Y., H. Taru, Y. Jin and C. C. Quinn, 2015 SYD-1C, UNC-40 (DCC) and SAX-3 (Robo) function interdependently to promote axon guidance by regulating the MIG-2 GTPase. PLoS Genet 11: e1005185.

40. Yamauchi, K., M. Yamazaki, M. Abe, K. Sakimura, H. Lickert et al., 2017 Netrin-1 Derived from the Ventricular Zone, but not the Floor Plate, Directs Hindbrain Commissural Axons to the Ventral Midline. Sci Rep 7: 11992.

41. Yan, J., D. L. Chao, S. Toba, K. Koyasako, T. Yasunaga et al., 2013 Kinesin-1 regulates dendrite microtubule polarity in Caenorhabditis elegans. Elife 2: e00133.

